# Scaling the PBWT for Long-Range Shared Ancestry Detection in Large Haplotype Panels

**DOI:** 10.64898/2025.12.01.691644

**Authors:** Uwaise Ibna Islam, Davide Cozzi, Travis Gagie, Rahul Varki, Vincenza Colonna, Erik Garrison, Paola Bonizzoni, Christina Boucher

**Affiliations:** Department of Computer and Information Science and Engineering,University of Florida,Gainesville,, FL, USA; Department of Informatics, Systems and Communication,University of Milano–Bicocca,Milan, 20126,, Italy; Faculty of Computer Science,Dalhousie University,Halifax,, NS, Canada; Department of Genetics, Genomics and Informatics,University of Tennessee Health Science Center (UTHSC),Memphis,, TN, USA; Institute of Genetics and Biophysics,National Research Council,Naples, 80111,, Italy; Department of Pediatrics,University of Tennessee Health Science Center (UTHSC),Memphis,, TN, USA

**Keywords:** Positional Burrows–Wheeler Transform (PBWT), set-maximal exact matches (SMEMs), Identity-by-Descent (IBD), Population Genomics, Pangenomics, Run-length encoding, Phased haploytpes, Shared ancestry

## Abstract

Detecting long shared ancestry tracts in large haplotype panels is central to IBD analysis, imputation, and local ancestry inference, and can be approximated computationally by finding Set-Maximal Exact Matches (SMEMs) between sequences. The Positional Burrows–Wheeler Transform (PBWT) provides an efficient index for these panels, yet current methods often enumerate all SMEMs, producing a large number of short, uninformative matches. We introduce Positional Boyer– Moore–Li (`PBML`), which restricts enumeration to SMEMs occurring in at least *k* haplotypes and spanning at least *L* sites (*kL*-SMEMs). `PBML` is the first algorithm for computing *KL*-SMEMs on top of a single compressed run-length encoded PBWT index reusable for any (*k, L*) without rebuilding. On the 1000 Genomes Project, `PBML` achieves 4.6× faster query time than *µ*-PBWT and 2.4× over Durbin’s PBWT with lower memory, scaling to 15.9× over *µ*-PBWT at 16 threads. On a 10,000-haplotype panel from the Tennessee BIG Initiative, a diverse admixed cohort, `PBML` outperforms *µ*-PBWT by up to 4.7× in *k*-SMEM finding. By applying both thresholds during traversal, `PBML` extracts biologically informative, population-shared segments while filtering millions of short matches, a capability not available in current tools. On the BIG panel, in about 10 seconds `PBML` finds 2,441 long tracts at (*k* = 50, *L* = 5000) shared by an average of 60 haplotypes against 1000 queries, significantly reducing the 4.8 million unfiltered SMEMs shared on average by 2 haplotypes. These results establish `PBML` as a scalable tool for targeted long-range shared ancestry detection in large, diverse panels.

## Introduction

By preserving phase at the chromosome scale, haplotype-resolved assemblies allow for precise mapping of admixture tracts [24, 7], elucidation of disease genetics [22, 13], estimation of homozygosity [23], and inference of local ancestry [36]. In line with this, several large-scale efforts are generating haplotype-resolved assemblies, including the UK Biobank [18], the Mexican Biobank [32], TOPMed [33], and All of Us [1]. To index and search these large panels efficiently, the Positional Burrows–Wheeler Transform (PBWT) provides a phase-aware representation. Originally proposed by Durbin [12], the PBWT stores a set of *h* sequences with *w* variant sites as an *h* × *w* binary matrix *S*[1 … *h*][1 … *w*], with rows maintained in co-lexicographic order. For a given haplotype panel, it supports efficient querying and haplotype matching in *O*(*hw*) time, and has been applied to haplotype phasing [6, 19], haplotype estimation [11], genotype imputation [30], genome database search [28], ancestry inference [35], and Identity-By-Descent (IBD) segment detection [25, 37, 14, 32]. Reflecting the central role of the PBWT in haplotype indexing and matching, recent work has focused on compressing the structure, extending its functionality, and accelerating support for various query types [28, 27, 26, 34].

A unifying formulation underlying many of these applications is the detection of shared inheritance between individuals in a PBWT panel. This can be cast as the problem of finding all set maximal exact matches (SMEMs) between a query sequence and the haplotype panel, where a SMEM is the longest subsequence shared between the query and at least one sequence represented in the PBWT. The seminal PBWT includes an algorithm for locating SMEMs [12], and subsequent work has optimized SMEM discovery using run-length encoding, sampling, and parallelism to achieve fast, memory-efficient haplotype matching [9, 31]. A remaining challenge is to restrict the search space, as enumerating all SMEMs often yields an extremely large number of short matches that inflate downstream analysis [15, 5]. To address this, previous work has been extended to consider SMEMs that occur at least *k* times in the panel, termed *k*-SMEMs [3].

Here, we further focus on *k*-SMEMs of length at least *L*, which we call *kL*-SMEMs: matches that occur in at least *k* panel haplotypes and span at least *L* sites. This restriction greatly reduces the number of reported matches while retaining those that are most useful in practice. To solve this problem, we introduce `PBML` (Positional Boyer-Moore-Li), a space- and time-efficient algorithm to find all *kL*-SMEMs. Conceptually, `PBML` builds on the Boyer-Moore-Li (BML) algorithm, which combines Li’s forward–backward MEM-finding strategy with Boyer-Moore-style skipping via Longest Common Prefix (LCP) and Longest Common Suffix (LCS) queries on the PBWT index [20, 4, 15, 21]. In particular, `PBML` skips right along the query haplotype and then extends matches leftward on run-length encoded forward and reverse PBWTs, enabling efficient length-constrained *k*-SMEM enumeration. Crucially, a single prebuilt index supports queries at any combination of *k* and *L* without rebuilding, enabling rapid exploration of the (*k, L*) parameter space.

We evaluated `PBML` on the 1000 Genomes Project (1KGP, 5,008 haplotypes) and on a 10,000 haplotype panel from the Tennessee BIG Initiative [8], a diverse Mid-South cohort with substantial African American representation. On 1KGP, we benchmark against existing PBWT-based methods and show that `PBML` achieves the fastest query time with the lowest memory among compressed indexes, with strong multi-threaded scaling due to a shared read-only index. On BIG, we demonstrate that a single prebuilt `PBML` index dominates *µ*-PBWT [3] across a sweep of *k* values in k-SMEM finding, avoiding the repeated index construction that *µ*-PBWT [3] requires for each threshold. We then show that increasing the minimum SMEM length *L* provides output-sensitive speedups while retaining over 95% site coverage at moderate thresholds, and that combining both parameters yields biologically targeted output: simultaneous (*k, L*) filtering isolates long, recurrently shared tracts characteristic of IBD segments, reducing millions of uninformative matches to a compact set of high-confidence candidates in seconds. The source code for `PBML` is publicly available at https://github.com/uwaiseibna/PBML.git.

## Preliminaries

### Representation of haplotypes

We represent phased haplotype data as a binary matrix *S* ∈ {0, 1}^*h×w*^, where each of the *h* rows corresponds to a haplotype and each of the *w* columns corresponds to a biallelic variant site ordered along the genome. An entry *S*[*i, j*] is equal to 0 if the haplotype *i* carries the reference allele at the site *j*, and 1 if it carries the alternate allele. For each column *j*, we write 0_*j*_ for the number of 0-bits in column *j*.

We note that we use 0-based indexing throughout, so rows are indexed 0, …, *h*−1 and columns 0, …, *w*−1.

### Forward PBWT

PBWT [12] is a data structure for storing and querying a haplotype panel *S* ∈ {0, 1}^*h×w*^ . For each column *j* ∈ {0, …, *w*−1}, the PBWT maintains a permutation *a*_*j*_ of {0, …, *h* − 1}, called the *prefix array*, which orders the haplotypes by the co-lexicographic order of their prefixes up to column *j* − 1. Formally, *a*_*j*_ [*i*] is the index of the *i*th haplotype in the ordering obtained by sorting the prefixes *S*[0, 0..*j* − 1], *S*[1, 0..*j* − 1], …, *S*[*h* − 1, 0..*j* − 1] in co-lexicographic order, i.e., lexicographic order when comparing prefixes from right to left (from position *j*−1 down to 0). Because each column refines the previous ordering via a stable partition according to the current column, haplotypes sharing a common prefix remain contiguous as *j* increases.

The PBWT matrix *P* ∈ {0, 1}^*h×w*^ is defined by permuting each column of *S* according to the corresponding prefix array. Specifically, at column 0 the prefix is empty, so *a*_0_ is the identity permutation and *P*[*i*, 0] = *S*[*i*, 0]. For *j* ≥ 1, *P*[*i, j*] = *S*[*a*_*j*_ [*i*], *j*]. We refer to this as the forward PBWT to distinguish it from the reverse PBWT defined below.

### Reverse PBWT

The reverse PBWT is defined analogously by processing the panel right-to-left (equivalently, by applying the PBWT to the column-reversed panel). It maintains, for each column *j*, a permutation 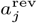 that orders haplotypes by the co-lexicographic order of their suffixes starting at column *j* +1. The reverse PBWT matrix *P* ^rev^ ∈ {0, 1}^*h×w*^ is defined by permuting each column of *S* according to 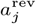, with *P* ^rev^[*i, w*− 1] = *S*[*i, w* − 1] and, for 0 ≤ *j* ≤ *w* − 2, 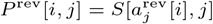.

### RLE-PBWT

A *run* in column *j* of *P* is a maximal block of consecutive identical bits in that column. We write *r*_*j*_ for the number of runs in column *j* and *r* = ∑_*j*_ *r*_*j*_ for the total number of runs in all columns. Thus, we refer to run-length encoding (RLE) as the representation of a PBWT column as a sequence of its runs, storing for each run its length and starting bit of each column.

### PBWT intervals

As a consequence of stable refinement as *j* increases, haplotypes sharing a common prefix (or suffix, in the reverse PBWT) occupy a contiguous interval [*s, e*] in the prefix array. Given a query *Q* ∈ {0, 1}^*w*^ and a starting column *c*, the set of haplotypes matching with a match length *ℓ, Q*[*c*..*c* + *ℓ* − 1] can be computed by iteratively refining an initial interval [0, *h* − 1] while scanning successive columns of the PBWT.

### LCP and LCS

Both *Longest Common Prefix (LCP)* and *Longest Common Suffix (LCS)* queries operate directly on the run-length–encoded PBWT columns, where each column *j* stores its starting bit, the run lengths, and the number of zeros 0_*j*_ . Let *Q*[0..*w* − 1] be the query haplotype. During a Longest Common Prefix query starting at column *c*, we maintain a PBWT interval [*s, e*] containing exactly the haplotypes that match the current query prefix, initialized to [0, *h* − 1]. At each column, we use rank queries to count how many 1-bits occur before and within this interval; these counts determine how the interval is mapped into the next column, either into the block of 0s or the block of 1s, depending on the query bit. The extension stops when fewer than *k* haplotypes remain in the interval or when we reach the query boundary (column *w* − 1 for LCP, column 0 for LCS). The LCP query returns the match length *ℓ*, the final interval [*s, e*], and the haplotype index *a*_*j*_ [*e*] at the upper boundary of this interval, tracked throughout the extension for later retrieval of all matching haplotypes.

### SMEMs

Given a panel *S* ∈ {0, 1}^*h×w*^ and a query *Q* ∈ {0, 1}^*w*^, we define a SMEM is a substring *Q*[*i*..*j*] with 0 ≤ *i* ≤ *j < w* such that: (1) there exists at least one haplotype *s* ∈ *S* such that *s*[*i*..*j*] = *Q*[*i*..*j*], and (2) the match is maximal with respect to the extension in either direction, i.e., no *s* ∈ *S* satisfies *s*[*i* − 1..*j*] = *Q*[*i* − 1..*j*] when *i* > 0, and no *s* ∈ *S* satisfies *s*[*i*..*j* + 1] = *Q*[*i*..*j* + 1] when *j < w* − 1.

Next, we define a *k-SMEM* a SMEM that occurs at least *k* haplotypes in *S* contain the match, and a *kL-SMEM* as a *k*-SMEM of length at least *L*.

## Methods

This section describes how `PBML` computes *kL*-SMEMs on the RLE-PBWT, and how we recover matching haplotypes efficiently. In particular, we first describe how to perform LCP and LCS queries directly on RLE-PBWT, which form the core primitive for matching a query haplotype against the panel. We then show how to combine them to enumerate all *k*-SMEMs of length ≥ *L* with Boyer-Moore-Li. After computing the interval of matching haplotypes, we give an algorithm to recover them. Lastly, we give the complexity analysis of `PBML` .

### LCP and LCS Queries on the RLE-PBWT

Both LCP and LCS queries operate on RLE columns, where each column *j* stores its starting bit, run lengths, and zero-count (i.e., 0_*j*_). We let *Q*[0 . . *w*−1] be the query haplotype. During an LCP query starting at column *c*, we maintain a PBWT interval [*s, e*] representing haplotypes matching the current prefix. The interval [*s, e*] is initialized to [0, *h*−1]. Then at each column *j*, we compute rank values *r*_*s*_ = rank_1_(*s*−1, *j*) and *r*_*e*_ = rank_1_(*e, j*), and update the interval based on query bit *Q*[*j*]:

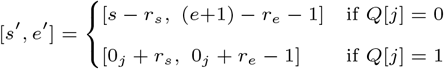

The extension ends when fewer than *k* haplotypes remain (i.e., *s*^*′*^ + *k* > *e*^*′*^ + 1) or the query boundary is reached, i.e., column *w* − 1 for LCP, and column 0 for LCS. LCP returns the match length *ℓ*, the final interval [*s, e*], and the haplotype index *a*_*j*_ [*e*] at the boundary of the final interval, tracked during extension for subsequent haplotype retrieval.

LCS queries apply the same interval-narrowing procedure on the reverse RLE-PBWT, processing columns from *c* toward 0, returning only the match length *ℓ* since haplotype retrieval only requires the RLE forward PBWT.

Both queries run in *O*(*r*)-time. Full algorithmic details appear in Supplementary Section B.

### Boyer–Moore–Li in the PBWT

Given the query haplotype *Q*[0..*w* − 1] and integers *L* and *k*, we now describe how to compute all *kL*-SMEMs between the PBWT and the query haplotype *Q*. We note that any SMEM of length at least *L* whose start lies in *Q*[0..*L* − 1] must include position *Q*[*L* − 1]. Consequently, we initiate the search at *Q*[*L* − 1] and match from right-to-left using an LCS query on the reverse RLE-PBWT, updating the interval at each step until fewer than *k* haplotypes remain at position *Q*[*i*]. At this point, there are two cases depending on the position *i* at which the match ends.

If *i* = 0, the backward extension reaches the beginning of *Q*. We then extend the current interval forward from *Q*[0] via an LCP query on the forward RLE-PBWT, which proceeds through column *p*−1 before fewer than *k* haplotypes remain at column *p*, producing the first SMEM *Q*[0..*p*−1]. To find the next SMEM, the algorithm checks whether any SMEM of length ≥ *L* covers position *p*. It performs a backward extension from *Q*[*p*] of length *ℓ*. If *ℓ* ≥ *L*, the corresponding forward extension yields the next SMEM. If *ℓ < L*, no SMEM of length ≥ *L* can cover *Q*[*p*], and moreover the first *L* − *ℓ* positions starting at *p* cannot begin such an SMEM either. The algorithm skips past them, analogous to the shift rule in Boyer–Moore, and continues from position *Q*[*p* + *L* − *ℓ*].

Otherwise, we know that *i* ≥ 1 and it follows that there does not exist any SMEM of length at least *L* begins in *Q*[0..*i*−1]. We therefore recurse on the suffix *Q*[*i*..*w*−1], starting at *Q*[*i*+*L*−1].

The recursion ends once the remaining suffix has length less than *L*. We note that when *L* = 1, the procedure simplifies to the standard forward-backward algorithm. We give the pseudocode in Supplementary Section A.

### Identifying Matching Haplotypes

To obtain the haplotypes in the interval [*s, e*] produced by an LCP query, we require access to the prefix array at the corresponding columns. However, storing the complete prefix array for all columns is memory-prohibitive for large panels. Instead, `PBML` adapts the Toehold Lemma and *ϕ* predecessor operation from the *r*-index [16, 29] to recover the haplotypes, which removes the necessity of storing the full prefix arrays.

We first show how to recover the haplotypes using *ϕ*. Given PBWT column *j, ϕ* query on *i*th haplotype of prefix array *a*_*j*_ [*i*], returns the previous haplotype in the prefix array, *a*_*j*_ [*i* − 1]; *ϕ*(*a*_*j*_ [*i*]) = *a*_*j*_ [*i* − 1]. During an LCP query starting at column *c*, we maintain the haplotype *a*_*j*_ [*e*] at the interval endpoint *e* as the interval narrows across successive columns. When extending from column *j* to column *j*+1, if the bit at position *e* in column *j* matches the query bit *Q*[*j*] then the same haplotype remains at the updated endpoint *e*^*′*^ in column *j* + 1. Otherwise, the interval endpoint moves to the previous run in the PBWT column, and we update the haplotype to the one at the end of the preceding run via a *ϕ* query. By the end of the LCP query, we have identified the haplotype at the final interval endpoint without accessing the full prefix array.

Given this tracked haplotype and the final interval [*s, e*], we enumerate all matching haplotypes by repeatedly applying *ϕ*: starting from *a*_*j*_ [*e*], we compute its co-lexicographic predecessor in the prefix array, then the predecessor of that, and so on, until we have traversed the entire interval. This recovers all *e* − *s* + 1 haplotype identities without materializing the prefix array.

To support *ϕ* efficiently, `PBML` maintains two auxiliary structures. The first is a per-haplotype *successor structure* that stores, for each haplotype, the PBWT columns where it starts a run, the haplotype immediately preceding it in the prefix array at each such column, and a pointer into that predecessor’s successor structure indicating the closest later column where the predecessor also began a run. The second is a per-run *run-sampled structure* that stores each run’s starting row offset, the haplotype index at the run start, and a pointer into the corresponding successor structure. Each column additionally stores the index of its first run for direct access. Full details are provided in Supplementary Section C.

### Complexity Analysis

We analyze PBML for a single query against a panel of *h* haplotypes over *w* sites. Let *r* denote the total number of runs across the forward and reverse run-length encoded PBWTs. Because `PBML` operates directly on the RLE-PBWT, both the index size and query cost depend on the number of runs rather than the full panel size. In large biobank datasets, long shared haplotype segments often make *r* much smaller than *hw*, enabling compact storage and efficient rank queries. During matching, `PBML` performs forward and backward extensions using LCP and LCS queries.

The efficiency of `PBML` is summarized by the following theorem. We give the proof of this Theorem in Supplementary Section D.

**Theorem 1** *Given integers k and L, a query Q* ∈ {0, 1}^*w*^, *and a panel S of h haplotypes over w sites*, `PBML` *finds all kL-SMEMs in S in O*(*r*)*-space and O*(*N*_vis_*r* + occ)*-time, where N*_vis_ *is the total number of columns visited, and* occ *is the number of kL-SMEMs occurrences in S*.

In practice, the threshold *L* allows `PBML` to skip positions that cannot begin a valid match, reducing the number of visited columns and improving runtime, especially when long matches are sparse.

## Results

### Experimental setup

`PBML` is implemented in `C++17` using SDSL [17] for bit vectors, OpenMP for multi-threaded queries, and htslib [2] for VCF/BCF parsing. The experiments were run on an AMD EPYC 75F3 32-Core Processor with 503 GB RAM, Red Hat Enterprise Linux 9, and g++ 11.5.0. Running time and peak memory were measured with `/usr/bin/time -v`. We compared `PBML` to *µ*-PBWT [9], dynamic *µ*-PBWT [31], and `PBWT` ^orig^ [12] by recording the construction time, peak memory, and query time.

### Datasets

We evaluated on two datasets. The first consists of panels from the 1,000 Genomes Project (1KGP) Phase 3: 2,504 individuals (5,008 haplotypes), filtered to biallelic SNPs using `bcftools` [10], with 1 to 6 million variants per chromosome. For each experiment, 1,000 haplotypes serve as queries and the remaining 4,008 as the reference panel.

The second dataset consists of the BIG data [8], a multi-institutional resource from a diverse mid-south population with substantial African American representation from Memphis: 5,500 individuals (11,000 haplotypes), phased with ShapeIT5 [19] with variant sites 0.9 million to 4.4 million per chromosome. Similarly to the 1KGP dataset, we use 1,000 haplotypes as queries and the remaining 10,000 haplotypes as the reference panel.

### Enumerating SMEMs in the 1KGP Data

Figure 2 summarizes single-threaded performance across all 22 autosomes. Build and query times are normalized per million variant sites (Table inside Figure 2); per-chromosome trends are shown in Figures 2 and Supplementary section F (multi-thread). `PBWT` ^orig^ achieves the fastest construction (1.6× faster than `PBML`), but is not compressed. Among compressed methods, `PBML` constructs 1.4 times faster than *µ*-PBWT and 5 times faster than dynamic *µ*-PBWT.

**Fig. 1.**
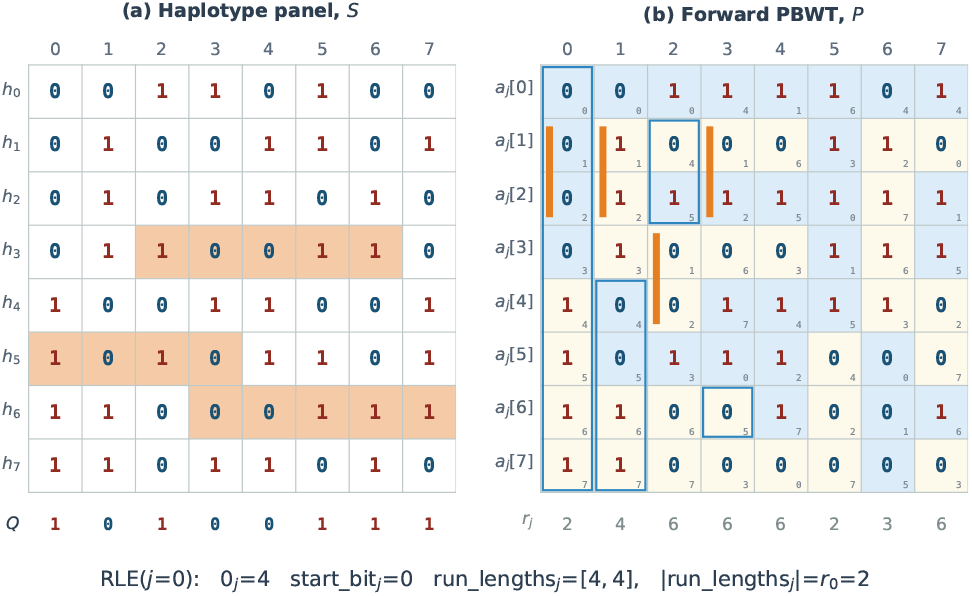
Worked example (*h* = 8, *w* = 8). **(a)** *S* with query *Q* and SMEMs shaded. **(b)** Forward PBWT *P*, rows ordered by *a*_*j*_ (subscripts show *a*_*j*_ [*i*]), runs shaded with counts *r*_*j*_ below, blue boxes tracing LCP interval [*s, e*] from column 0, orange bars tracking PBWT interval: {*h*_1_, *h*_2_} across columns, and RLE representation of column 0 at bottom.

**Fig. 2.**
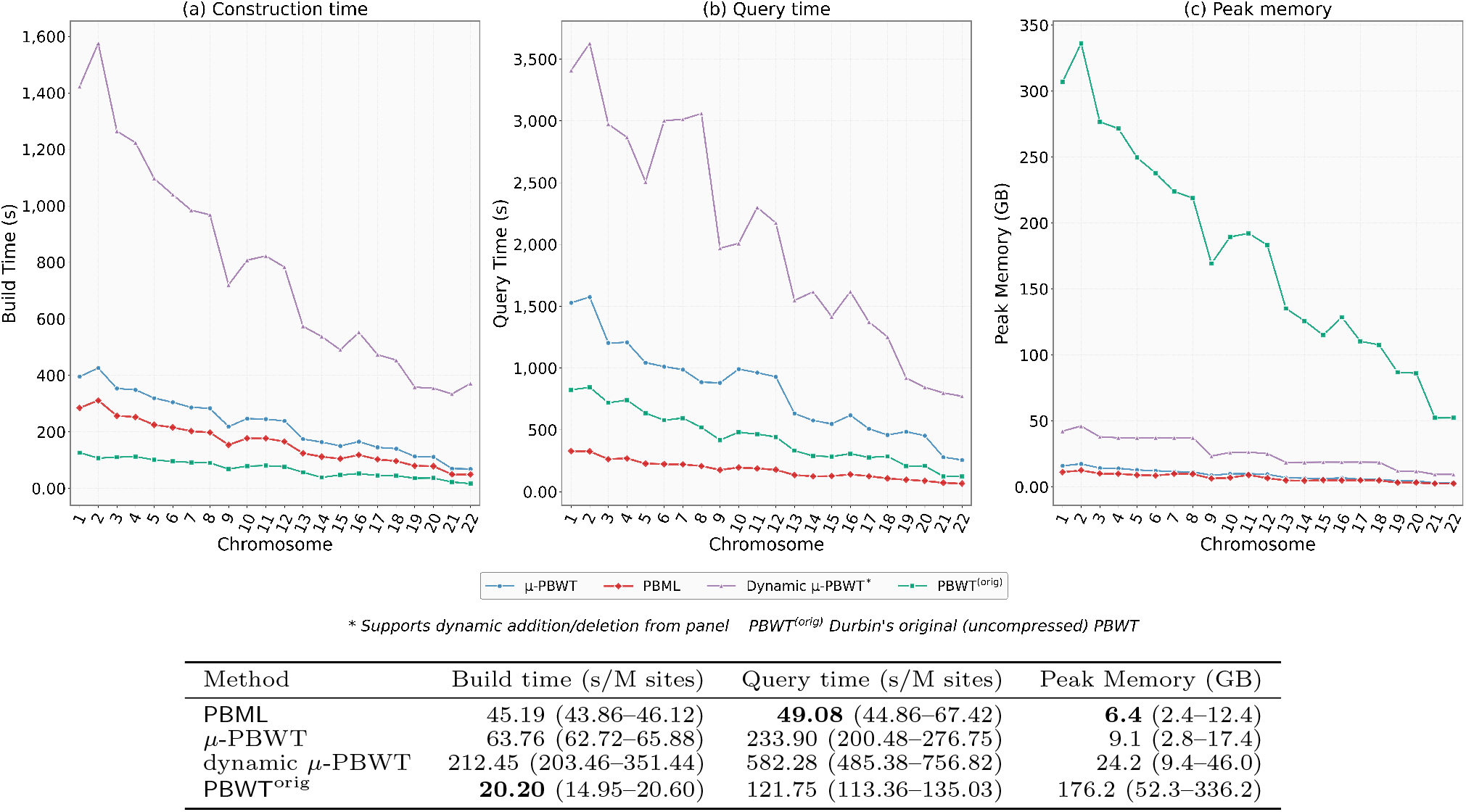
Single-threaded performance across all 22 autosomes of the 1000 Genomes Project (1KGP) panel. Per-chromosome (a) index construction time, (b) total query time for 1,000 haplotypes, and (c) peak memory usage are shown for `PBML`, *µ*-PBWT, dynamic *µ*-PBWT, and `PBWT` ^orig^ (times normalized per million variant sites). The table summarizes median (min–max) build time, query time, and peak memory across chromosomes (1KGP, *L* = 1, 1,000 queries), with best values in **bold**.

For 1,000 queries, `PBML` achieves the fastest query time, which is 4.6× faster than *µ*-PBWT and 2.4× faster than `PBWT`^orig^. This advantage stems from the forward-backward algorithm, which resolves each SMEM through a single LCS-LCP pair rather than scanning the full query to compute matching statistics, and from Boyer-Moore skipping, which avoids revisiting sites already covered by a previously reported SMEM. Moreover, the peak memory of `PBML` is 23% lower than *µ*-PBWT and 96% lower than `PBWT`^orig^.

We analyze multi-thread (16 threads) performance in Supplementary section F (`PBWT` ^orig^ and dynamic *µ*-PBWT are excluded because they do not support multi-threading). `PBML` achieves 8.2× speedup with near-constant memory (an average increase of 0.6%), because all threads share a single read-only index and allocate only thread-local output buffers. In contrast, *µ*-PBWT achieves a 2.4× speedup with a 15.9% memory increase due to per-thread matching statistics. At 16 threads, `PBML` queries 15.9× faster than *µ*-PBWT while using 1.5× less memory.

### Enumerating *k*-SMEMs in the BIG Dataset

In biobank data, many SMEMs that occur only once or a few times correspond to private or near-private mutations and provide little value for downstream analyses. Hence, requiring each SMEM to occur in at least *k* panel haplotypes filters these rare matches and retains only those supported by recurrent sharing across the cohort. We evaluate this on BIG chromosome 22 (10,000 haplotypes, 893,511 sites, 1,000 queries, single-threaded). Here, we set *L* = 1, allowing *k*-SMEMs to have arbitrary length.

Table 1 shows that `PBML` is faster for every value of *k*, the advantage increasing from 1.2× to 4.7× as *k* increases from 1 to 100. In this range, the runtime of *µ*-PBWT [3] increases by a factor of 6.2, while `PBML` increases by a factor of 1.7. A key design advantage is index reuse: `PBML` builds the index once and then answers queries repeatedly, whereas *µ*-PBWT [3] rebuilds the index for each *k*, accumulating over 3,500 seconds of redundant construction time. In addition, the memory usage of `PBML` remains constant at 2.5GB for all values of *k* greater than 1. The slightly higher usage when *k* equals 1 (2.8 GB) reflects the per-haplotype output format used in the default case, whereas for all other values of *k* we output a single line per SMEM with all matching haplotypes listed. In contrast, the memory usage of *µ*-PBWT [3] increases from 3.8 GB to 11.1 GB when *k* = 100 for identical output format.

**Table 1.** *k*-SMEM query performance on BIG chromosome 22. `PBML` uses a prebuilt index (141.5 seconds build, mean 3.8 seconds load time); whereas *µ*-PBWT [3] rebuilds index from panel per *k*, incurring a mean build time of 759.7 seconds.

| $k$ | PBML | | μ-PBWT | |
| --- | --- | --- | --- | --- |
|  | Time (s) | Mem (GB) | Time (s) | Mem (GB) |
| 1 | **590** | **2.8** | 736 | 3.8 |
| 5 | **827** | **2.5** | 1,531 | 4.4 |
| 10 | **861** | **2.5** | 2,067 | 4.9 |
| 50 | **912** | **2.5** | 3,524 | 7.6 |
| 100 | **979** | **2.5** | 4,572 | 11.1 |

### Investigating the Effect of *L*

Next, we evaluate the effect of the minimum SMEM length threshold *L* on chromosome 22 for both the 1KGP and BIG panels at *k* = 1 (Figure 3). We define *coverage* as the fraction of variant sites in a query haplotype spanned by at least one reported SMEM; when aggregating multiple queries, we take the union of all SMEMs before computing this fraction.

**Fig. 3.**
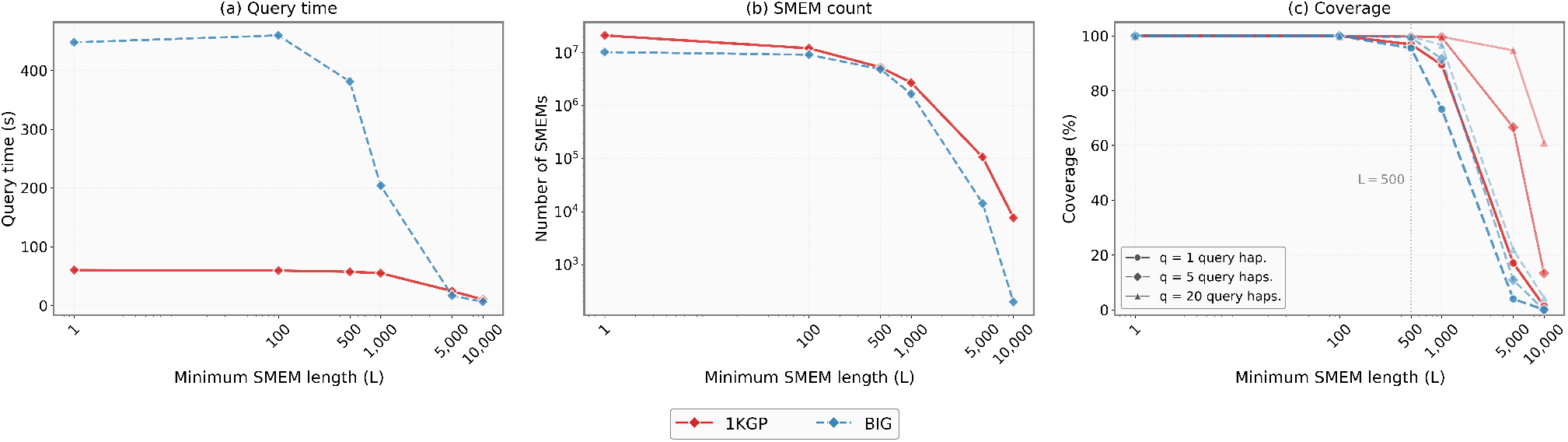
Effect of minimum SMEM length threshold (*L*) on chromosome 22 for the 1KGP (4,008 haplotypes, solid) and BIG (10,000 haplotypes, dashed) panels. (a) Query time decreases with *L* as fewer SMEMs are enumerated. (b) SMEM count on log scale. (c) Coverage (fraction of query sites spanned by ≥1 SMEM) for 1, 5, and 20 aggregated query haplotypes; the vertical line marks *L* = 500.

In 1KGP, increasing *L* reduces the number of SMEMs, from 21.1 million at *L* = 1 to 5.3 million at *L* = 500, and further to 7,674 at *L* = 10, 000. The query time decreases accordingly, from 60.3 seconds to 10.2 seconds (about an 83% reduction). The coverage at *L* = 500 is 96.9% for a single query and 99.8% when aggregating five. Increasing *L* reduces query time and maintains high coverage at moderate *L* values (Supplementary section H).

In BIG, the effect is more pronounced due to the larger panel and the higher density of short matches. The SMEM count decreases from 10.2 million in *L* = 1 to 4.9 million in *L* = 500 and to 14,283 in *L* = 5, 000, yielding a 37× reduction in the query time (590 to 16 seconds). At *L* = 500, BIG retains 95.5% single-query coverage and 99.4% when aggregating five queries, despite a 53% reduction in SMEM count. Compared to 1KGP, coverage degrades more steeply at the same *L* because the larger\ panel generates more short matches whose removal impacts site coverage more substantially.

In both panels, moderate values of *L* (e.g., 500 to 1,000) provide a favorable tradeoff between coverage and computational savings. However, *L* alone cannot distinguish long private matches from long shared matches; combining *L* with a minimum frequency threshold *k* addresses this limitation.

### Enumerating *kL*-SMEMs in the BIG Dataset

The preceding sections evaluate *k* and *L* independently and show that increasing *k* excludes rare SMEMs while increasing *L* excludes short ones. Yet neither parameter alone identifies SMEMs that are both long and widely shared. For example, setting *k* = 1 and *L* = 5, 000 on chromosome 22 yields 8,983 long SMEMs (mean length 6,179), but each is shared by only 1.6 haplotypes on average, likely uninformative for downstream analyses such as IBD detection. Conversely, *k* = 50 with *L* = 1 produces 9.8M SMEMs shared across 91.4 haplotypes on average, but with a mean length of only 440.

`PBML` applies both thresholds simultaneously on a single reusable prebuilt index. Figure 4 evaluates all nine (*k, L*) combinations with *k* ∈ {1, 10, 50} and *L* ∈ {1, 500, 5000} across all 22 autosomes of the BIG panel. Panel 4(a) plots mean SMEM length against mean haplotypes per SMEM for each chromosome under each (*k, L*) configuration. The nine configurations separate into distinct clusters that reveal the complementary roles of the two parameters: increasing *L* shifts clusters rightward (longer SMEMs) while increasing *k* shifts them upward (more widely shared). Only the (*k* = 50, *L* = 5, 000) cluster occupies the upper-right region, corresponding to long, recurrently shared tracts characteristic of IBD segments. On chromosome 22, this configuration yields 2,441 SMEMs with a mean length of 5,815 shared by 58.5 haplotypes, compared with 4.8 million short, mostly private SMEMs at (*k* = 1, *L* = 1). The tight within-cluster grouping of all 22 chromosomes confirms that these filtering characteristics are consistent genome-wide.

**Fig. 4.**
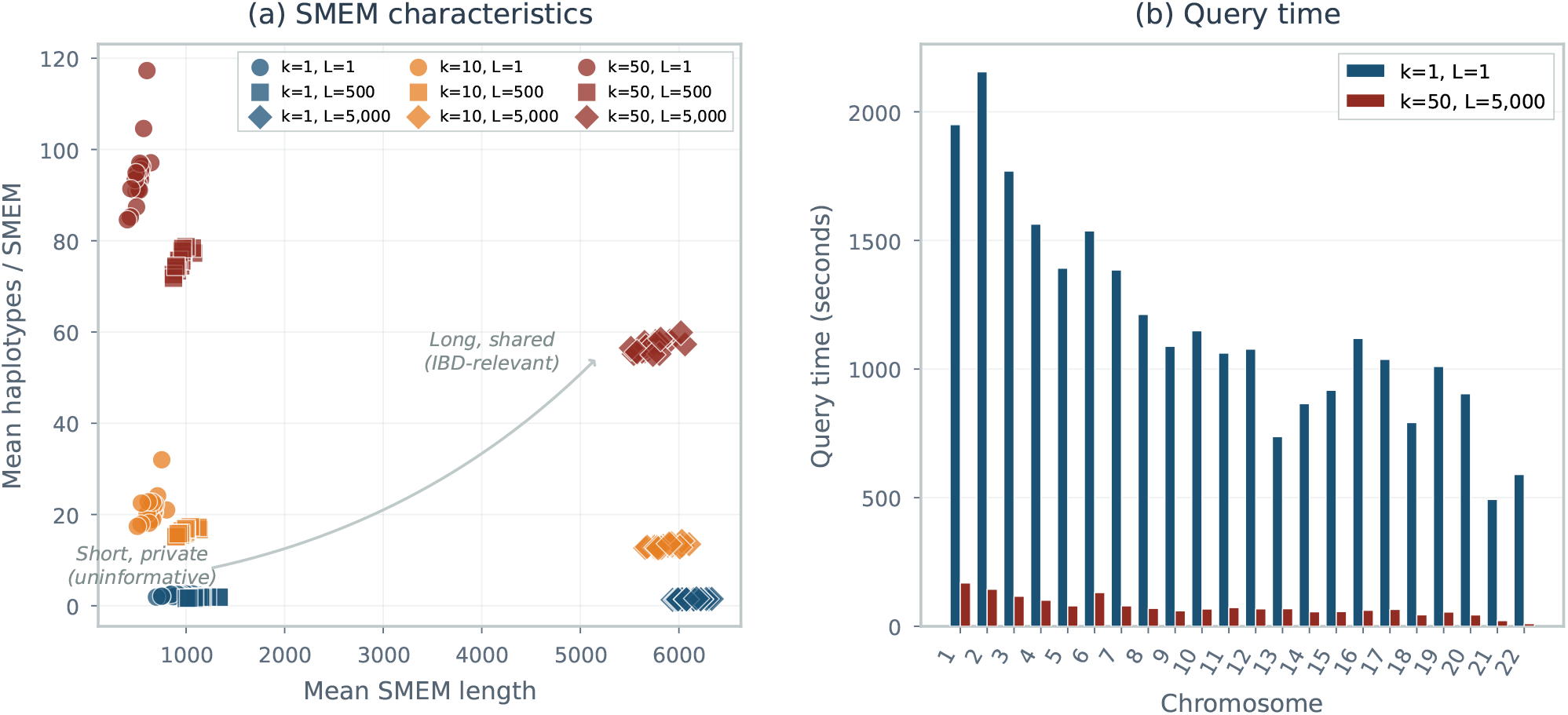
*kL*-SMEMs across all BIG autosomes (10,000 haplotypes, 1,000 queries, single-threaded). (a) Each point represents one chromosome under one (*k, L*) configuration, plotted by mean SMEM length (*x*-axis) and mean haplotypes per SMEM (*y*-axis). (b) Per-chromosome query time comparing the unfiltered baseline (*k* = 1, *L* = 1) with the IBD-focused configuration (*k* = 50, *L* = 5, 000).

Panel (b) shows the corresponding query times. IBD-focused threshold (*k* = 50, *L* = 5, 000) reduces the genome-wide query time from 25,806 seconds (∼7.2 hours) to 1,642 seconds (∼27 minutes), a 15.7× speedup driven by a 1, 050× reduction in output size (221.5M to 211K SMEMs). Per-chromosome speedups range from 10.8× to 59.0× (median 16.0×). The full per-chromosome characterization of haplotype sharing across all configurations is provided in Supplementary section G.

## Conclusion

By building a single *O*(*r*)-space index that supports queries in any combination of *k* and *L* without rebuilding, `PBML` shifts the SMEM enumeration from exhaustive listing to targeted extraction: the user specifies what constitutes a biologically meaningful match and the algorithm delivers only those. This is particularly relevant for applications such as IBD detection and haplotype imputation, where short or private matches are noise and the segments of interest are both long and recurrently shared across the panel. No prior method supports both of these parameterizations. Future work includes validating detected segments against recombination maps through integration with local-ancestry inference pipelines, and extending the algorithm to multi-allelic and graph-based PBWT representations to support structurally complex variation.

## Acknowledgements

We gratefully acknowledge the Tennessee Biorepository and Integrative Genomics (BIG) Initiative for providing access to the biobank data from the Mid-South population. We thank the participants, clinical staff, and investigators whose efforts made these data available.

## Notes

### Competing Interest Statement

The authors have declared no competing interest.

### Summary of Updates

We have significantly updated the PBML framework to enhance its scalability and biological utility for large-scale genomic studies. The core improvements are as follows: Support for kL-SMEMs: The algorithm has been extended to support dual-threshold filtering. PBML now identifies Set-Maximal Exact Matches (SMEMs) that occur in at least k haplotypes and span at least L genomic sites. This allows researchers to focus specifically on matches that are both long and frequently shared within a population, effectively filtering out millions of short or rare matches that often clutter traditional PBWT outputs. Full Run-Length Encoding (RLE) Integration: To optimize memory efficiency and performance on large panels, we have transitioned all query-time data structures to a run-length encoded format. PBML no longer relies on SDSL bit vectors for the PBWT representation. Instead, we utilize RLE-based logic to handle Longest Common Prefix (LCP) and Longest Common Successor (LCS) queries. This change reduces the memory footprint and allows the tool to scale more effectively with the number of haplotypes. Persistent Indexing: We have added comprehensive index support to the implementation. The PBWT index can now be constructed and saved to disk, enabling a "build once, query later" workflow. This eliminates the computational overhead of rebuilding the index for different k and L parameters or during iterative querying sessions. Enhanced Benchmarking: Our performance evaluation has been updated to include comparisons against the latest dynamic mu-PBWT implementation. These benchmarks demonstrate that PBML achieves up to 4.6x faster query times on the 1000 Genomes Project and up to 15.9x speedup when utilizing 16 threads. Tests on the Tennessee BIG Initiative panel further confirm that PBML outperforms dynamic mu-PBWT by up to 4.7x in k-SMEM finding tasks while maintaining a significantly lower memory profile. These updates establish PBML as a robust, production-ready tool for analyzing shared ancestry in diverse, biobank-scale haplotype collections.

