## Supplementary material for "Scaling the PBWT for Long-Range Shared Ancestry Detection in Large Haplotype Panels": supp

### Supplement

#### A. Pseudocode for PBML

**Algorithm 1** PBML querying algorithm. RLE-PBWTs$^{(*)}$ denotes the forward and reverse PBWTs stored in run-length encoded form.

```
Require: Query Q[0..w − 1], minimum length L, minimum occurrences k, RLE-PBWTs(*)
Ensure: All kL-SMEMs of Q against S
left ← 0
while left + L − 1 < w do
    col ← left + L − 1
    ℓ ← LCS(Q, col, k)                                  ▷ reverse RLE-PBWT
    if ℓ < L then
        left ← left + max(1, L − ℓ)                     ▷ Boyer–Moore skip
        continue
    end if
    start ← col − ℓ + 1
    (ℓ′, [s, e], a_j[e]) ← LCP(Q, start, k)             ▷ forward RLE-PBWT
    end ← start + ℓ′ − 1
    if ℓ′ ≥ L then
        retrieve haplotype IDs in [s, e] from a_j[e] via φ
        report SMEM Q[start..end]
    end if
    left ← end − L + 2
end while
```

#### B. RLE-PBWT rank queries and LCS/LCP algorithms

For each column $j$ of the RLE-PBWT, the index stores the bit value of the first run `startBit`$[j] \in \{0, 1\}$, an array of run lengths `runLens`$[j]$, a pointer `colPtr`$[j]$ into the global run-length array, and the count of 0-bits $0_j$ used for interval updates. Here, $r_j =$ `colPtr`$[j{+}1] -$ `colPtr`$[j]$ is number of runs in column $j$.

Given a position $i$ in column $j$, we compute $\mathrm{rank}_1(i, j)$ by scanning runs until reaching position $i$:

**Algorithm 2** RLE rank query

```
Require: Position i, column j, RLE-PBWT
Ensure: rank_1(i, j) and bit value at position i
rank ← 0; pos ← 0; bit ← startBit[j]
for p ← colPtr[j] to colPtr[j + 1] − 1 do
    len ← runLens[p]
    if pos + len > i then
        if bit = 1 then
            rank ← rank + (i − pos + 1)
        end if
        return (rank, bit)
    end if
    if bit = 1 then
        rank ← rank + len
    end if
    pos ← pos + len
    bit ← 1 − bit
end for
```

This runs in $O(r_j)$ time and $O(1)$ space. In practice, we compute $\mathrm{rank}_1(s{-}1, j)$ and $\mathrm{rank}_1(e, j)$ in a single scan for interval updates.

##### *LCS and LCP algorithms*

Both algorithms perform PBWT interval narrowing: starting from the full range $[0, h{-}1]$, they iteratively restrict the interval to haplotypes matching the query. LCS extends backward using the reverse RLE-PBWT; LCP extends forward using the forward RLE-PBWT. LCP additionally tracks $a_j[e]$, the original haplotype at the interval end, initialized from `end_prefs`$[c] = a_c[h{-}1]$ (precomputed during construction) and updated via $\phi$ operations when the interval end crosses a run boundary. $\textsc{FindRun}(e, j)$

locates the run containing position $e$ in column $j$ via binary search over cumulative run boundaries. The structures `run_phi_info` and `hap_succ` are the run-sampled and successor structures described in the Methods.

**Algorithm 3** LCS: Longest common suffix query

**Require:** Query $Q$, starting column $c$, min occurrences $k$, reverse RLE-PBWT
**Ensure:** Maximum backward extension length $\ell$

```
s ← 0; e ← h − 1; ℓ ← 0
while c − ℓ ≥ 0 do
    j ← c − ℓ
    (r_s, r_e) ← RankPair(s−1, e, j)                ▷ O(r_j) via Alg. 2
    if Q[j] = 0 then
        s′ ← s − r_s; e′ ← (e + 1) − r_e
    else
        s′ ← 0_j + r_s; e′ ← 0_j + r_e
    end if
    if s′ + k > e′ then
        break                                        ▷ fewer than k matches
    end if
    s ← s′; e ← e′ − 1; ℓ ← ℓ + 1
end while
return ℓ
```

**Algorithm 4** LCP: Longest common prefix query

**Require:** Query $Q$, starting column $c$, min occurrences $k$, forward RLE-PBWT
**Ensure:** Extension length $\ell$, final interval $[s, e]$, interval-ending haplotype $a_j[e]$

```
s ← 0; e ← h − 1; ℓ ← 0
a_j[e] ← end_prefs[c]
while c + ℓ < w do
    j ← c + ℓ
    (r_s, r_e, b_e) ← RankAndBit(s−1, e, j)         ▷ rank pair + bit at e
    if Q[j] = 0 then
        s′ ← s − r_s; e′ ← (e + 1) − r_e
    else
        s′ ← 0_j + r_s; e′ ← 0_j + r_e
    end if
    if s′ + k > e′ then
        break
    end if
    if b_e ≠ Q[j] then                               ▷ interval end crosses run boundary
        run ← FindRun(e, j)                          ▷ O(log r_j)
        (hap, idx) ← run_phi_info[run]
        a_j[e] ← hap_succ[hap].preds[idx]
    end if
    s ← s′; e ← e′ − 1; ℓ ← ℓ + 1
end while
return (ℓ, [s, e], a_j[e])
```

LCS performs $\ell$ iterations and LCP performs $\ell'$ iterations, each requiring an $O(r_j)$ rank query and an $O(1)$ interval update; LCP additionally performs an $O(\log r_j)$ run lookup when the interval-ending haplotype changes runs. The total cost per extension is $O(\sum r_j)$ over the columns visited. In uncompressed PBWT, rank queries take $O(1)$ with precomputed rank arrays but require $O(h \times w)$ space; the RLE approach trades $O(1) \to O(r_j)$ query time for $O(h \times w) \to O(r)$ space, a favorable exchange since $r_j \ll h$ in genomic data with high haplotype similarity.

#### C. Supporting $\phi$ on the RLE-PBWT

Nishimoto and Tabei [1] showed how to support $\phi$ on the $r$-index in constant worst-case time through run-splitting, and Zakeri et al. [2] demonstrated that splitting is unnecessary in practice, providing an efficient table-lookup-based implementation. We adapt the latter approach to the PBWT.

Given the interval-ending haplotype $a_j[e]$ produced by an LCP query, the first $\phi$ operation requires locating this haplotype's position within its successor structure via binary search over the $n_s$ columns where that haplotype starts a run. This takes $O(\log n_s)$

time. Once located, the successor entry directly provides the predecessor haplotype $a_j[e-1]$ and a hint pointer into the predecessor's own successor structure, indicating the closest later column where the predecessor also began a run. Each subsequent $\phi$ operation follows these hint pointers, requiring only a constant-time table lookup. Thus, retrieving all $e - s + 1$ haplotypes in the interval costs $O(\log n_s + (e - s))$: one binary search followed by $e - s$ constant-time steps.

#### D. Proof of Theorem 1

We restate Theorem 1: PBML finds all $kL$-SMEMs in $O(r)$-space and time $O(N_{\text{vis}} r + \text{occ})$.

*Proof* **Space.** Both the forward and reverse PBWTs are stored in run-length encoded form, requiring $O(r)$ space, where $r$ is the total number of runs across all columns. The $\phi$-supporting structures (successor arrays and run-sampled structures for haplotype recovery) record entries strictly at run boundaries, ensuring their total size is also bounded by $O(r)$. The per-column metadata (starting bit, zero-count, column pointer, first-run index, end-preference) requires $O(w)$ words. Since each of the $w$ columns contains at least one run, we strictly have $r \geq w$, the metadata space is asymptotically absorbed into $O(r)$. Therefore, the total index size is $O(r)$.

**Time.** We analyze the total time complexity T by bounding the cost of three distinct phases: (i) SMEM-finding iterations, (ii) skipped iterations, and (iii) haplotype reporting.

In each iteration $i$ across the $\sigma$ SMEM-finding iterations, the algorithm performs a backward extension, visiting $lcs_i$ columns, and a forward extension, visiting $lcp_i$ columns. We recall that to perform those extension queries, each step from column $j$ to the adjacent column is bounded by the cost of a rank query: $O(r_j)$. Following a successful match, the search position advances by $lcp_i - lcs_i + 1$ columns. Since the successful iterations strictly tile the query $Q$ of length $w$, we have: $\sum_i^{\sigma}(lcp_i - lcs_i + 1) = w$. Hence, we can express the total number of forward column visits in terms of the backward ones: $\sum_{i=1}^{\sigma} lcp_i = w + \sum_{i=1}^{\sigma} lcs_i - \sigma$. Denoting $\Lambda = \sum_i^{\sigma} lcs_i - \sigma$ the total excess backward extension, we can express the total forward visits as $\sum_i^{\sigma} lcp_i = w + \Lambda$. Similarly, we can express the total backward visits as $\sum_i^{\sigma} lcs_i = \Lambda + \sigma$. Given these premises, the exact number of column visits across all the $\sigma$ SMEM-finding iterations is $N_{\text{vis}} = \sum_{i=1}^{\sigma}(lcs_i + lcp_i) = w + 2\Lambda + \sigma$. Consequently, the time required in the SMEM-finding iterations is bounded by $O\left(\sum_{t=1}^{N_{\text{vis}}} r_{j_t}\right)$, where $j_t$ is the index of the column visited at step $t$.

When $L > 1$, the LCS procedure may terminate with a maximum backward extension of length $\ell < L$. In such cases, the algorithm performs a Boyer–Moore skip, advancing the search position by $\max(1, L - \ell)$ columns. Each skipped iteration visits exactly $\ell \leq L - 1$ columns before advancing by at least one column. Because the total advancement of the search pointer across the entire execution is strictly bounded by the total number of columns $w$, the cumulative number of columns visited by all skipped iterations is bounded by $O(w)$. Since $r_j \geq 1$ for each column $j$, the cost of the skipped iterations is absorbed by the cost of the SMEM-finding iterations: $O\left(\sum_{t=1}^{N_{\text{vis}}} r_{j_t}\right)$.

Furthermore, for each of the $\sigma$ SMEM intervals, we report the corresponding haplotype subset. Locating the relative position of the interval-ending haplotype requires a binary search on its $n_{s_i}$-length successor array, taking $O(\log n_{s_i})$ time. The remaining haplotypes in the interval are then recovered via hint-guided $\phi$-operations. Each $\phi$-operation requires $O(1)$ amortized time. Hence, summing these costs over all the $\sigma$ SMEM intervals, retrieving all the occ haplotypes of the whole SMEM set takes $O\left(\sum_{i=1}^{\sigma} \log n_{s_i}\right)$ for the binary searches and $O(occ)$ for the remaining haplotypes.

Finally, we observe that each rank query costs $r_{j_t}$ in a single column; hence, each cost is trivially bounded by the total number of runs $r$. Therefore, the total time spent in the visited columns $O\left(\sum_{t=1}^{N_{\text{vis}}} r_{j_t}\right)$ is strictly bounded by $O(N_{\text{vis}} r)$. Furthermore, since each successor array size $n_{s_i}$ cannot exceed $r$, the cumulative cost of the $\sigma$ binary searches to locate the interval-ending haplotype $O\left(\sum_{i=1}^{\sigma} \log n_{s_i}\right)$ is at most $O(\sigma \log r)$. Recalling from our earlier derivation that $N_{\text{vis}} = w + 2\Lambda + \sigma$, where $w \geq 1$ and $\Lambda \geq 0$, it trivially follows that $\sigma \leq N_{\text{vis}}$ and that $O(\sigma \log r)$ is absorbed into $O(N_{\text{vis}} r)$.

The final time complexity is derived by adding to $O(N_{\text{vis}} r)$ the cost $O(occ)$ of locating all the remaining matching haplotypes. □

**Corollary 1** In the worst case, the total excess backward extension is $\Lambda = O(w^2)$. This scenario arises when the algorithm identifies $\sigma = O(w)$ heavily overlapping SMEMs, each of which advances the search position by only a single column, forcing the backward extension to revisit $O(w)$ columns per iteration. Consequently, the total number of column visits becomes $N_{\text{vis}} = O(w^2)$ and the overall worst-case time complexity evaluates to $O(w^2 \rho + \text{occ})$.

Remark 1 *In genomic data, high linkage disequilibrium keeps $\Lambda \ll w^2$. The Boyer–Moore skip logic (parameter $L$) further reduces both $\sigma$ and $\Lambda$ by bypassing positions that cannot begin a long SMEM, keeping the matching time near-linear in practice.*

#### E. Example: finding SMEMs with $L = 4$

We demonstrate PBML on the matrix $S'$ in Table 1 using the forward (Table 2a) and reverse PBWT (Table 2b), with minimum SMEM length $L = 4$ and $k = 1$. Successful LCS–LCP query pairs that produce SMEMs are highlighted in alternating yellow and cyan; failed LCS queries that do not meet the minimum length are highlighted in lavender.

We begin with *left* $= 0$ and perform an LCS query on the reverse PBWT (Table 2b) at column *col* $= 3$. Starting from the full interval $[0, 19]$, we narrow backward to column 0, giving $\ell = 4$. The subsequent LCP query on the forward PBWT (Table 2a) starts at column 0 with interval $[0, 19]$ and extends until fewer than $k$ haplotypes remain at column 6. The final valid interval $[11, 14]$ at column 5 contains haplotypes $\{8, 11, 12, 13\}$, yielding an SMEM of length 6 spanning across column 0 to column 5. We advance *left* to $5 - 4 + 2 = 3$.

**Haplotype panel**

| $h\backslash w$ | 0 | 1 | 2 | 3 | 4 | 5 | 6 | 7 | 8 | 9 | 10 | 11 | 12 | 13 | 14 |
|---|---|---|---|---|---|---|---|---|---|---|---|---|---|---|---|
| 0 | 1 | 0 | 0 | 1 | 0 | 0 | 0 | 0 | 0 | 0 | 0 | 1 | 1 | 0 | 1 |
| 1 | 1 | 0 | 0 | 1 | 1 | 0 | 0 | 1 | 0 | 0 | 0 | 0 | 0 | 1 | 1 |
| 2 | 1 | 0 | 0 | 1 | 1 | 0 | 0 | 1 | 0 | 0 | 0 | 1 | 0 | 0 | 1 |
| 3 | 1 | 0 | 0 | 1 | 1 | 0 | 0 | 1 | 0 | 0 | 0 | 1 | 0 | 0 | 1 |
| 4 | 0 | 1 | 0 | 1 | 0 | 1 | 0 | 0 | 0 | 0 | 0 | 1 | 0 | 0 | 1 |
| 5 | 0 | 1 | 0 | 1 | 0 | 1 | 0 | 0 | 0 | 0 | 0 | 1 | 0 | 0 | 1 |
| 6 | 0 | 1 | 0 | 1 | 0 | 1 | 0 | 0 | 0 | 0 | 0 | 1 | 0 | 0 | 1 |
| 7 | 0 | 1 | 0 | 1 | 0 | 1 | 0 | 0 | 0 | 0 | 0 | 0 | 1 | 0 | 1 |
| 8 | 0 | 1 | 0 | 0 | 1 | 0 | 0 | 0 | 0 | 1 | 1 | 1 | 0 | 0 | 1 |
| 9 | 0 | 1 | 0 | 1 | 0 | 0 | 0 | 0 | 1 | 0 | 0 | 0 | 0 | 1 | 1 |
| 10 | 0 | 1 | 0 | 1 | 0 | 0 | 0 | 0 | 1 | 0 | 0 | 0 | 0 | 1 | 1 |
| 11 | 0 | 1 | 0 | 0 | 1 | 0 | 0 | 0 | 0 | 0 | 1 | 1 | 0 | 0 | 0 |
| 12 | 0 | 1 | 0 | 0 | 1 | 0 | 0 | 0 | 1 | 0 | 1 | 1 | 0 | 0 | 1 |
| 13 | 0 | 1 | 0 | 0 | 1 | 0 | 0 | 0 | 1 | 0 | 1 | 1 | 0 | 0 | 1 |
| 14 | 0 | 1 | 0 | 0 | 0 | 0 | 0 | 0 | 1 | 0 | 0 | 0 | 1 | 0 | 1 |
| 15 | 0 | 1 | 0 | 0 | 0 | 0 | 0 | 0 | 1 | 0 | 0 | 0 | 1 | 0 | 1 |
| 16 | 0 | 1 | 0 | 1 | 0 | 0 | 0 | 0 | 0 | 0 | 0 | 1 | 1 | 0 | 1 |
| 17 | 1 | 1 | 0 | 0 | 0 | 1 | 0 | 0 | 0 | 0 | 0 | 1 | 1 | 0 | 1 |
| 18 | 0 | 1 | 1 | 0 | 1 | 0 | 0 | 0 | 0 | 0 | 0 | 1 | 0 | 0 | 1 |
| 19 | 0 | 1 | 1 | 0 | 1 | 0 | 1 | 0 | 0 | 0 | 0 | 0 | 1 | 0 | 1 |
| $Q$ | 0 | 1 | 0 | 0 | 1 | 0 | 1 | 0 | 0 | 0 | 1 | 1 | 1 | 0 | 1 |

**Table 1.** Example haplotype matrix $S'$ (20 haplotypes, 15 sites) with SMEMs for query $Q$ highlighted in yellow.

**(a) Forward PBWT**

| $S[a_j]/w$ | 0 | 1 | 2 | 3 | 4 | 5 | 6 | 7 | 8 | 9 | 10 | 11 | 12 | 13 | 14 |
|---|---|---|---|---|---|---|---|---|---|---|---|---|---|---|---|
| 0 | $1_{0}$ | $1_{4}$ | $0_{0}$ | $1_{0}$ | $1_{8}$ | $0_{14}$ | $0_{14}$ | $0_{14}$ | $1_{14}$ | $0_{0}$ | $0_{0}$ | $1_{0}$ | $1_{7}$ | $1_{1}$ | $1_{18}$ |
| 1 | $1_{1}$ | $1_{5}$ | $0_{1}$ | $1_{1}$ | $1_{11}$ | $0_{15}$ | $0_{15}$ | $0_{15}$ | $1_{15}$ | $0_{16}$ | $0_{16}$ | $1_{16}$ | $1_{19}$ | $1_{9}$ | $1_{4}$ |
| 2 | $1_{2}$ | $1_{6}$ | $0_{2}$ | $1_{2}$ | $1_{12}$ | $1_{17}$ | $0_{0}$ | $0_{0}$ | $0_{0}$ | $1_{8}$ | $1_{11}$ | $1_{18}$ | $0_{1}$ | $1_{10}$ | $1_{5}$ |
| 3 | $1_{3}$ | $1_{7}$ | $0_{3}$ | $1_{3}$ | $1_{13}$ | $0_{0}$ | $0_{9}$ | $0_{9}$ | $1_{9}$ | $0_{11}$ | $0_{18}$ | $1_{17}$ | $1_{14}$ | $0_{18}$ | $1_{6}$ |
| 4 | $0_{4}$ | $1_{8}$ | $0_{4}$ | $1_{4}$ | $0_{14}$ | $1_{4}$ | $0_{10}$ | $0_{10}$ | $1_{10}$ | $0_{18}$ | $0_{17}$ | $1_{4}$ | $1_{15}$ | $0_{4}$ | $1_{2}$ |
| 5 | $0_{5}$ | $1_{9}$ | $0_{5}$ | $1_{5}$ | $0_{15}$ | $1_{5}$ | $0_{16}$ | $0_{16}$ | $0_{16}$ | $0_{17}$ | $0_{4}$ | $1_{5}$ | $0_{9}$ | $0_{5}$ | $1_{3}$ |
| 6 | $0_{6}$ | $1_{10}$ | $0_{6}$ | $1_{6}$ | $0_{17}$ | $1_{6}$ | $0_{8}$ | $0_{8}$ | $0_{8}$ | $0_{4}$ | $0_{5}$ | $1_{6}$ | $0_{10}$ | $0_{6}$ | $0_{11}$ |
| 7 | $0_{7}$ | $1_{11}$ | $0_{7}$ | $1_{7}$ | $1_{18}$ | $1_{7}$ | $0_{11}$ | $0_{11}$ | $0_{11}$ | $0_{5}$ | $0_{6}$ | $0_{7}$ | $1_{0}$ | $0_{2}$ | $1_{12}$ |
| 8 | $0_{8}$ | $1_{12}$ | $0_{8}$ | $0_{8}$ | $1_{19}$ | $0_{9}$ | $0_{12}$ | $0_{12}$ | $1_{12}$ | $0_{6}$ | $0_{7}$ | $0_{19}$ | $1_{16}$ | $0_{3}$ | $1_{13}$ |
| 9 | $0_{9}$ | $1_{13}$ | $0_{9}$ | $1_{9}$ | $0_{0}$ | $0_{10}$ | $0_{13}$ | $0_{13}$ | $1_{13}$ | $0_{7}$ | $0_{19}$ | $0_{1}$ | $0_{18}$ | $0_{11}$ | $1_{8}$ |
| 10 | $0_{10}$ | $1_{14}$ | $0_{10}$ | $1_{10}$ | $1_{1}$ | $0_{16}$ | $0_{18}$ | $0_{18}$ | $0_{18}$ | $0_{19}$ | $0_{1}$ | $1_{2}$ | $1_{17}$ | $0_{12}$ | $1_{7}$ |
| 11 | $0_{11}$ | $1_{15}$ | $0_{11}$ | $0_{11}$ | $1_{2}$ | $0_{8}$ | $1_{19}$ | $1_{1}$ | $0_{17}$ | $0_{1}$ | $0_{2}$ | $1_{3}$ | $0_{4}$ | $0_{13}$ | $1_{19}$ |
| 12 | $0_{12}$ | $1_{16}$ | $0_{12}$ | $0_{12}$ | $1_{3}$ | $0_{11}$ | $0_{1}$ | $1_{2}$ | $0_{4}$ | $0_{2}$ | $0_{3}$ | $0_{14}$ | $0_{5}$ | $0_{8}$ | $1_{14}$ |
| 13 | $0_{13}$ | $1_{18}$ | $0_{13}$ | $0_{13}$ | $0_{4}$ | $0_{12}$ | $0_{2}$ | $1_{3}$ | $0_{5}$ | $0_{3}$ | $0_{14}$ | $0_{15}$ | $0_{6}$ | $0_{7}$ | $1_{15}$ |
| 14 | $0_{14}$ | $1_{19}$ | $0_{14}$ | $0_{14}$ | $0_{5}$ | $0_{13}$ | $0_{3}$ | $0_{17}$ | $0_{6}$ | $0_{14}$ | $0_{15}$ | $0_{9}$ | $0_{2}$ | $0_{19}$ | $1_{0}$ |
| 15 | $0_{15}$ | $0_{0}$ | $0_{15}$ | $0_{15}$ | $0_{6}$ | $0_{18}$ | $0_{17}$ | $0_{4}$ | $0_{7}$ | $0_{15}$ | $0_{9}$ | $0_{10}$ | $0_{3}$ | $0_{14}$ | $1_{16}$ |
| 16 | $0_{16}$ | $0_{1}$ | $0_{16}$ | $1_{16}$ | $0_{7}$ | $0_{19}$ | $0_{4}$ | $0_{5}$ | $0_{19}$ | $0_{9}$ | $0_{10}$ | $1_{11}$ | $0_{11}$ | $0_{15}$ | $1_{17}$ |
| 17 | $1_{17}$ | $0_{2}$ | $1_{18}$ | $0_{17}$ | $0_{9}$ | $0_{1}$ | $0_{5}$ | $0_{6}$ | $0_{1}$ | $0_{10}$ | $1_{12}$ | $1_{12}$ | $0_{12}$ | $0_{0}$ | $1_{1}$ |
| 18 | $0_{18}$ | $0_{3}$ | $1_{19}$ | $0_{18}$ | $0_{10}$ | $0_{2}$ | $0_{6}$ | $0_{7}$ | $0_{2}$ | $0_{12}$ | $1_{13}$ | $1_{13}$ | $0_{13}$ | $0_{16}$ | $1_{9}$ |
| 19 | $0_{19}$ | $1_{17}$ | $0_{17}$ | $0_{19}$ | $0_{16}$ | $0_{3}$ | $0_{7}$ | $0_{19}$ | $0_{3}$ | $0_{13}$ | $1_{8}$ | $1_{8}$ | $0_{8}$ | $0_{17}$ | $1_{10}$ |
| $Q$ | 0 | 1 | 0 | 0 | 1 | 0 | 1 | 0 | 0 | 0 | 1 | 1 | 1 | 0 | 1 |

**(b) Reverse PBWT**

| $S[a_j^{rev}]/w$ | 0 | 1 | 2 | 3 | 4 | 5 | 6 | 7 | 8 | 9 | 10 | 11 | 12 | 13 | 14 |
|---|---|---|---|---|---|---|---|---|---|---|---|---|---|---|---|
| 0 | $1_{0}$ | $1_{14}$ | $0_{14}$ | $1_{0}$ | $1_{18}$ | $1_{7}$ | $0_{7}$ | $1_{1}$ | $0_{1}$ | $0_{1}$ | $0_{1}$ | $1_{11}$ | $0_{11}$ | $0_{11}$ | $1_{0}$ |
| 1 | $1_{1}$ | $1_{15}$ | $0_{15}$ | $1_{16}$ | $0_{0}$ | $1_{4}$ | $1_{19}$ | $0_{7}$ | $1_{9}$ | $0_{9}$ | $0_{9}$ | $1_{2}$ | $1_{0}$ | $0_{0}$ | $1_{1}$ |
| 2 | $1_{2}$ | $1_{17}$ | $0_{17}$ | $1_{9}$ | $0_{16}$ | $1_{5}$ | $0_{4}$ | $0_{19}$ | $1_{10}$ | $0_{10}$ | $0_{10}$ | $1_{3}$ | $0_{2}$ | $1_{1}$ | $1_{2}$ |
| 3 | $1_{3}$ | $1_{11}$ | $1_{18}$ | $1_{10}$ | $1_{11}$ | $1_{6}$ | $0_{5}$ | $1_{2}$ | $0_{7}$ | $0_{7}$ | $0_{7}$ | $1_{4}$ | $0_{3}$ | $0_{2}$ | $1_{3}$ |
| 4 | $0_{14}$ | $1_{8}$ | $0_{11}$ | $0_{14}$ | $1_{8}$ | $0_{18}$ | $0_{6}$ | $1_{3}$ | $1_{14}$ | $0_{14}$ | $0_{14}$ | $1_{5}$ | $0_{4}$ | $0_{3}$ | $1_{4}$ |
| 5 | $0_{15}$ | $1_{12}$ | $0_{8}$ | $0_{15}$ | $0_{9}$ | $0_{0}$ | $0_{18}$ | $0_{4}$ | $1_{15}$ | $0_{15}$ | $0_{15}$ | $1_{6}$ | $0_{5}$ | $0_{4}$ | $1_{5}$ |
| 6 | $1_{17}$ | $1_{13}$ | $0_{12}$ | $1_{7}$ | $0_{10}$ | $0_{16}$ | $0_{0}$ | $0_{5}$ | $0_{19}$ | $0_{19}$ | $0_{19}$ | $1_{8}$ | $0_{6}$ | $0_{5}$ | $1_{6}$ |
| 7 | $0_{11}$ | $0_{0}$ | $0_{13}$ | $1_{4}$ | $0_{14}$ | $1_{17}$ | $0_{16}$ | $0_{6}$ | $0_{2}$ | $0_{2}$ | $1_{11}$ | $1_{12}$ | $1_{7}$ | $0_{6}$ | $1_{7}$ |
| 8 | $0_{8}$ | $1_{16}$ | $1_{19}$ | $1_{5}$ | $0_{15}$ | $0_{11}$ | $0_{17}$ | $0_{18}$ | $0_{3}$ | $0_{3}$ | $0_{2}$ | $1_{13}$ | $0_{8}$ | $0_{7}$ | $1_{8}$ |
| 9 | $0_{12}$ | $1_{9}$ | $0_{0}$ | $1_{6}$ | $1_{12}$ | $0_{8}$ | $0_{11}$ | $0_{0}$ | $0_{4}$ | $0_{4}$ | $0_{3}$ | $1_{18}$ | $0_{12}$ | $0_{8}$ | $1_{9}$ |
| 10 | $0_{13}$ | $1_{10}$ | $0_{16}$ | $0_{17}$ | $1_{13}$ | $0_{9}$ | $0_{8}$ | $0_{16}$ | $0_{5}$ | $0_{5}$ | $0_{4}$ | $0_{1}$ | $0_{13}$ | $1_{9}$ | $1_{10}$ |
| 11 | $0_{16}$ | $1_{7}$ | $0_{9}$ | $0_{18}$ | $1_{1}$ | $0_{10}$ | $0_{9}$ | $0_{17}$ | $0_{6}$ | $0_{6}$ | $0_{5}$ | $0_{9}$ | $1_{14}$ | $1_{10}$ | $0_{11}$ |
| 12 | $0_{9}$ | $1_{4}$ | $0_{10}$ | $0_{11}$ | $1_{2}$ | $0_{14}$ | $0_{10}$ | $0_{11}$ | $0_{18}$ | $0_{18}$ | $0_{6}$ | $0_{10}$ | $1_{15}$ | $0_{12}$ | $1_{12}$ |
| 13 | $0_{10}$ | $1_{5}$ | $0_{7}$ | $0_{8}$ | $1_{3}$ | $0_{15}$ | $0_{14}$ | $0_{8}$ | $0_{0}$ | $0_{0}$ | $1_{8}$ | $1_{0}$ | $1_{16}$ | $0_{13}$ | $1_{13}$ |
| 14 | $0_{7}$ | $1_{6}$ | $0_{4}$ | $0_{12}$ | $1_{19}$ | $0_{12}$ | $0_{15}$ | $0_{9}$ | $0_{16}$ | $0_{16}$ | $1_{12}$ | $0_{7}$ | $1_{17}$ | $0_{14}$ | $1_{14}$ |
| 15 | $0_{4}$ | $0_{1}$ | $0_{5}$ | $0_{13}$ | $0_{7}$ | $0_{13}$ | $0_{12}$ | $0_{10}$ | $0_{17}$ | $0_{17}$ | $1_{13}$ | $0_{14}$ | $0_{18}$ | $0_{15}$ | $1_{15}$ |
| 16 | $0_{5}$ | $0_{2}$ | $0_{6}$ | $1_{1}$ | $0_{4}$ | $0_{1}$ | $0_{13}$ | $0_{14}$ | $0_{11}$ | $0_{11}$ | $0_{18}$ | $0_{15}$ | $1_{19}$ | $0_{16}$ | $1_{16}$ |
| 17 | $0_{6}$ | $0_{3}$ | $0_{1}$ | $1_{2}$ | $0_{5}$ | $0_{2}$ | $0_{1}$ | $0_{15}$ | $1_{12}$ | $1_{8}$ | $0_{0}$ | $1_{16}$ | $0_{1}$ | $0_{17}$ | $1_{17}$ |
| 18 | $0_{18}$ | $1_{18}$ | $0_{2}$ | $1_{3}$ | $0_{6}$ | $0_{3}$ | $0_{2}$ | $0_{12}$ | $1_{13}$ | $0_{12}$ | $0_{16}$ | $1_{17}$ | $0_{9}$ | $0_{18}$ | $1_{18}$ |
| 19 | $0_{19}$ | $1_{19}$ | $0_{3}$ | $0_{19}$ | $0_{17}$ | $0_{19}$ | $0_{3}$ | $0_{13}$ | $0_{8}$ | $0_{13}$ | $0_{17}$ | $0_{19}$ | $0_{10}$ | $0_{19}$ | $1_{19}$ |
| $Q$ | 0 | 1 | 0 | 0 | 1 | 0 | 1 | 0 | 0 | 0 | 1 | 1 | 1 | 0 | 1 |

**Table 2.** Forward and reverse PBWTs on $S'$ from Table 1. Each cell shows $b_{a_j[i]}$ (resp. $b_{a_j^{\text{rev}}[i]}$). Yellow/cyan mark alternating LCS–LCP interval pairs; lavender marks failed LCS queries ($\ell < L$); blended colors indicate overlap.

With *left* $= 3$, LCS at column *col* $= 6$ extends backward to column 3 ($\ell = 4$). LCP from column 3 extends until column 10; the final valid interval $[10, 10]$ at column 9 contains haplotype $\{19\}$, giving an SMEM of length 7 spanning across column 3 to column 9. We advance to *left* $= 9 - 4 + 2 = 7$.

With *left* $= 7$, LCS at column *col* $= 10$ extends backward to column 7 ($\ell = 4$). LCP from column 7 extends until column 12; the final valid interval $[16, 16]$ at column 11 contains haplotype $\{11\}$, giving an SMEM of length 5 spanning across column 7 to column 11. We advance to *left* $= 11 - 4 + 2 = 9$.

With *left* $= 9$, LCS at column *col* $= 12$ yields $\ell = 2 < L$. The Boyer–Moore skip advances *left* by $\max(1, L - \ell) = 2$ to *left* $= 11$; no LCP query is performed.

With *left* $= 11$, LCS at column *col* $= 14$ extends backward to column 11 ($\ell = 4$). LCP from column 11 extends to column 14, reaching the end of the query. The final interval $[14, 16]$ contains haplotypes $\{0, 16, 17\}$, giving an SMEM of length 4 spanning across column 11 to column 14. This terminates the procedure.

#### F. Multi-thread Performance

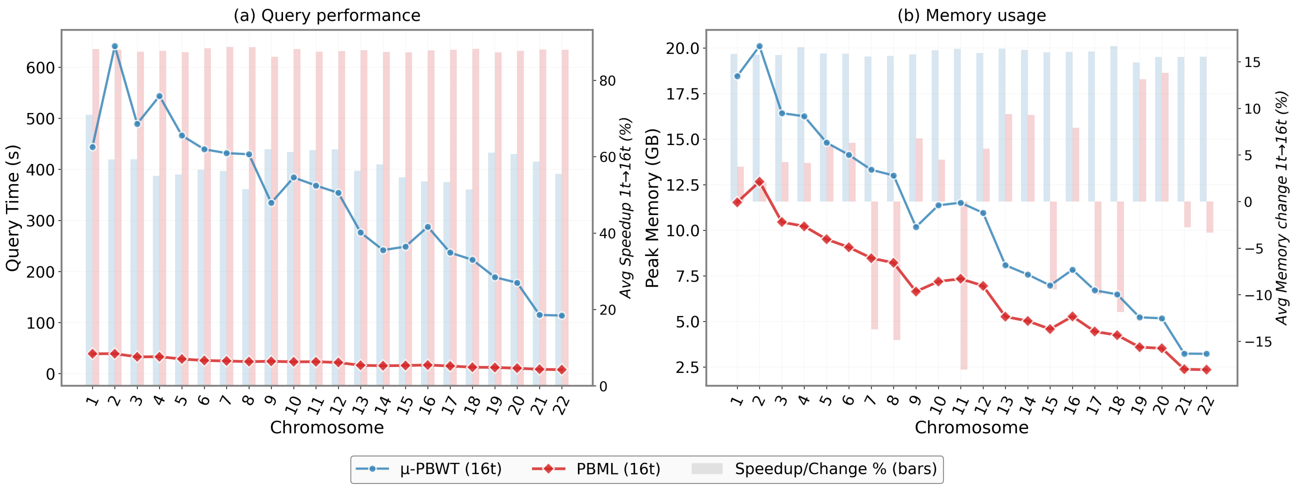


**Fig. 1. Multi-threaded comparison (16 threads) across all chromosomes for PBML and $\mu$-PBWT.** PBWT$^{orig}$ and dynamic $\mu$-PBWT are excluded as they do not support multi-threading. Dual y-axes show absolute values (left, solid lines) and relative changes from single-threaded baseline (right, semi-transparent bars).

#### G. Genome-wide $kL$-SMEM Haplotype Sharing

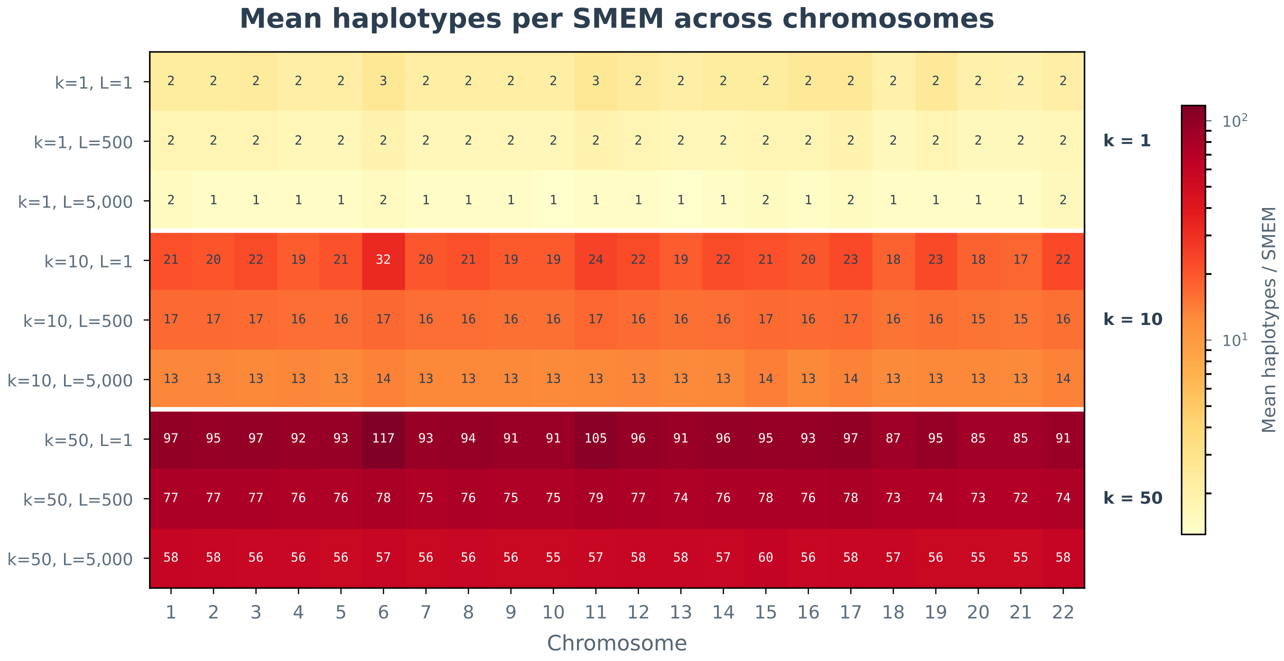


**Fig. 2. Mean haplotypes per SMEM across all BIG autosomes for each $(k, L)$ configuration.** Rows correspond to $(k, L)$ pairs grouped by $k$; columns correspond to chromosomes 1 to 22. Within each $k$ group, haplotype sharing is stable across values of $L$ and across chromosomes, confirming that the frequency parameter $k$ governs sharing while $L$ controls match length independently.

Figure 4 in the main text shows that the nine $(k, L)$ configurations form distinct clusters in SMEM length versus haplotype sharing space, and that joint filtering at $(k = 50,\ L = 5,000)$ yields consistent speedups across all 22 autosomes. Supplementary Figure **??** provides the complete per-chromosome detail underlying those results. The heatmap confirms two patterns: first, haplotype sharing is governed primarily by $k$, not $L$, as within each $k$ group the per-chromosome values are nearly uniform across all three values of $L$; and second, for a given $(k, L)$ pair, variation across chromosomes is small despite a roughly $5\times$ range in the number of variant sites (0.9 to 4.4 million). These observations confirm that the filtering behavior reported on chromosome 22 generalizes across the genome.

#### H. Genome-wide $L$-scaling on 1KGP

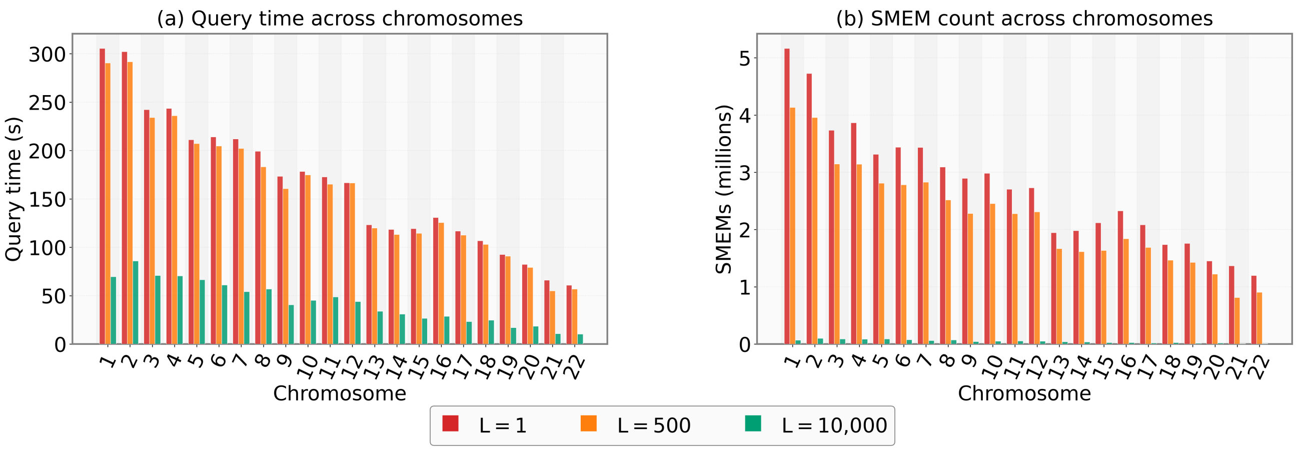


**Fig. 3.** Genome-wide performance of PBML on 1KGP at three minimum SMEM length thresholds ($L = 1$, 500, 10,000) and $k = 1$ across all 22 autosomes. (a) Query time per chromosome. (b) SMEM count per chromosome (millions). At $L$=500, total query time decreases modestly (3,487 s vs. 3,637 s at $L$=1) with a 19% reduction in SMEM count. At $L$=10,000, total runtime drops by 74.2% (937 s) with a 98.3% reduction in SMEM count. The per-chromosome speedup is consistent (median 74.6%), demonstrating predictable output-sensitive acceleration.
